# Nutrient stoichiometry shapes microbial coevolution

**DOI:** 10.1101/183657

**Authors:** Megan L. Larsen, Steven W. Wilhelm, Jay T. Lennon

**Affiliations:** Department of Biology, Indiana University, Bloomington, IN 47405; Department of Microbiology, University of Tennessee, Knoxville, TN 37996

**Keywords:** bacteria, phage, resources, resistance, arms race, networks

## Abstract

Coevolution is a force contributing to the generation and maintenance of biodiversity. It is influenced by environmental conditions including the scarcity of essential resources, which can drive the evolution of defense and virulence traits. We conducted a long-term chemostat experiment where the marine cyanobacterium *Synechococcus* was challenged with a lytic phage under nitrogen (N) or phosphorus (P) limitation. This manipulation of nutrient stoichiometry altered the stability of host-parasite interactions and the underlying mode of coevolution. By assessing infectivity with >18,000 pairwise challenges, we documented directional selection for increased phage resistance, consistent with arms-race dynamics while phage infectivity fluctuated through time, as expected when coevolution is driven by negative frequency-dependent selection. The resulting infection networks were 50 % less modular under N-versus P-limitation reflecting host-range contraction and asymmetric coevolutionary trajectories. Nutrient stoichiometry affects eco-evolutionary feedbacks in ways that may alter the dynamics and functioning of environmental and host-associated microbial communities.

## INTRODUCTION

In order to grow and reproduce, organisms must assimilate resources from the environment to meet their nutritional and energetic demands. Ecological stoichiometry is a theoretical framework that explicitly considers the mass balance of materials and energy in the environment and the individuals that incorporate them (Sterner & Elser 2002). Of the approximately 25 elements contained in biomass, nitrogen (N) and phosphorus (P) are two of the most limiting and essential nutrients. N and P are needed for the synthesis of major macromolecules, including nucleic acids, ribosomes, proteins, and cellular membranes that collectively influence an organism’s performance. However, the degree to which N and P are regulated differs among major groups of taxa. For example, the biomass stoichiometry of primary producers tends to be flexible, reflecting the supply of nutrients in their environment (Sterner & Elser 2002). In contrast, the biomass stoichiometry of consumer populations (e.g., viruses, invertebrates, mammals) is homeostatically regulated and therefore tends to remain constant even when the nutrient content of their resources fluctuates (Sterner & Elser 2002). Such differences can create a nutritional imbalance between primary producers and consumers, which has profound consequences for a wide range of ecological processes including resource competition, host-parasite dynamics, and ecosystem functioning (Smith & Holt 1996; Hall 2009; Aalto *et al*. 2015).

Nutrient stoichiometry can also influence evolutionary processes. Nutrient limitation often leads to a reduction in population size, which can diminish the efficiency of selection and result in the accumulation of deleterious mutations through genetic drift (Kimura 1962). However, many populations have adaptations that allow individuals to contend with absolute and relative resource scarcity. For example, the disproportionate use of nucleotides that vary in N content reduces stoichiometric mismatch between an organism and its environment (Elser *et al*. 2011). Similarly, natural selection can operate on the material costs of gene expression in ways that lead to the sparing of elements that are normally used in highly expressed proteins (Bragg & Wagner 2009). These genetic responses to nutrient limitation may give rise to the evolution of organismal stoichiometry. For example, long-term carbon limitation led to increased N and P content of experimentally evolved bacterial populations (Turner *et al*. 2017). However, evolutionary responses to nutrient stoichiometry may depend on the identity of the limiting nutrient (Gresham *et al*. 2008). While N limitation can select for stress-related or catabolic genes with lower guanine-cytosine content (Acquisti *et al*. 2009), P limitation can favor the replacement of phospholipids with sulfolipids in the cell membranes of certain microorganisms (Van Mooy *et al*. 2006).

When considering multiple interacting species, the combined effects of nutrient stoichiometry may give rise to eco-evolutionary feedbacks. Such feedbacks occur when ecological interactions affect evolutionary processes, which in turn modify species interactions and ecological dynamics (Haloin & Strauss 2008). For example, the rapid evolution of functional traits can produce diminished oscillations, longer periods of cycling, and phase-shifted population densities between hosts and their parasites, a phenomenon referred to as “cryptic dynamics” (Yoshida *et al*. 2007). Theory suggests that cryptic dynamics can arise when nutrient stoichiometry alters the stability of antagonistic species interactions (Yamamichi *et al*. 2015), which may ultimately intensify the modes of coevolution, namely arms-race dynamics and negative frequency-dependent selection (Aalto *et al*. 2015). However, links among nutrient stoichiometry, eco-evolutionary feedbacks, and coevolution remain to be tested.

Major advances in evolutionary ecology have been made through experimental studies of microbial communities. In particular, bacteria and phage are ideal for studying eco-evolutionary feedbacks owing to their large population sizes, rapid growth rates, and experimental tractability. Moreover, bacteria and phage dynamics are critical for understanding the structure and function of microbial food webs, especially in aquatic ecosystems (Suttle 2007). For example, *Synechococcus* is a diverse and widely distributed group of primary producers in the world ocean with an estimated global abundance of 10^26^ cells (Flombaum *et al*. 2013). While *Synechococcus* must contend with both N- and P-limitation, it is also subject to a high degree of phage-induced mortality, which may lead to nutrient-dependent coevolutionary dynamics. These coevolutionary processes can create feedbacks that not only influence population dynamics, but also ecosystem functioning, including the turnover of N and P (Lennon & Martiny 2008).

In this study, we tested how nutrient stoichiometry affects host-parasite eco-evolutionary dynamics in a simplified community where a single genotype of marine *Synechococcus* was infected with a phage strain in continuous culture using chemostats that were supplied with either N- or P-limited media. We isolated hundreds of host and phage strains, which were challenged against one another to document phenotypic changes in resistance and infectivity over the course of the experiment. With this information, we conducted network analyses that allowed us to quantify the degree of nestedness and modularity, which can serve as signatures of shed light on the underlying modes of coevolution, namely arms race dynamics and negative frequency dependent selection (Weitz *et al*. 2013). Our findings reveal that nutrient stoichiometry is a bottom-up force that regulates eco-evolutionary feedbacks in ways that alter coevolutionary processes.

## METHODS

### Strains and media

We evaluated the effects of nutrient stoichiometry on the eco-evolutionary dynamics of the marine cyanobacterium *Synechococcus* WH7803 and a lytic T4-like phage belonging to the Myoviridae family of phage (S-RIM8). Changes to nutrient supply can alter the equilibrium density of microbial populations in continuous culture (i.e., chemostats), which in turn, may affect cyanobacteria-phage contact rates (Rhee 1978). Therefore, we adjusted both the concentrations and ratios of N and P in modified AN artificial seawater medium (Tables S1-S3) to induce N- or P-limitation while maintaining similar equilibrium densities of *Synechococcus* between the stoichiometry treatments prior to phage addition. Specifically, the N-limited medium had an N : P ratio of 10 : 1 (KNO_3_; N = 220 µM) while the P-limited medium had a N : P ratio of 40 : 1 (K_2_HPO_4_; *P* = 11 µM). Growth assays confirmed that *Synechococcus* was limited by N under low N : P supply and by P under high N : P supply (Fig. S1).

### Nutrient limitation assay

We tested for nutrient limitation in *Synechococcus* by measuring growth rates on cells that had been maintained under low (10 : 1) and high (40 : 1) supply ratio of N and P. We acclimated *Synechococcus* semi-continuously under constant light (20 µE m^−1^ s^−1^) at 25 °C for three serial transfers in either N- or P-limited medium (n = 10). We then inoculated each replicate cell line into fresh medium that doubled the concentration of N (N-limited = 440 µM, P-limited = 880 µM) or P (N-limited = 44 µM, P-limited = 22 µM). Next, we monitored *Synechococcus* population densities for seven days *via* autofluorescence (ex: 550 nm, em: 570 nm) with a Biotek Synergy Mx plate reader (Winooski, VT, USA). With these data, we estimated the maximum growth rate for each culture using a modified Gompertz equation (Lennon et al. 2007) and calculated the percent change in growth rate with additional N or P as compared to the control, which contained no additional N or P.

### Chemostat experiment

We supplied ten chemostats, each with a 40 mL operating volume, with N-limited or P-limited medium at a dilution rate of 1 d^−1^ (Tables S1-S3). The chemostats were maintained in a Percival growth chamber (Perry, IA, USA) at 25 °C on a 14:10 light : dark cycle under 20 µE m^−2^ s^−1^ and homogenized with magnetic stir bars following inoculation with a single-colony strain of *Synechococcus* WH7803. We allowed the *Synechococcus* populations to equilibrate in the chemostats prior to initiating the “phage-amended” treatment by introducing an aliquot of a plaque-purified S-RIM8 to three randomly chosen chemostats in each nutrient treatment to achieve a multiplicity of infection (phage : cyanobacteria ratio) of approximately 10. To document the potential influence of stoichiometry on population dynamics and population size, we maintained “no phage” chemostats with only *Synechococcus* in both stoichiometry treatments (Fig. S2).

### Community dynamics

We tracked *Synechococcus* and phage densities in each chemostat via epifluorescent microscopy every other day for 172 days. *Synechococcus* populations were enumerated by concentrating samples from a chemostat onto 0.22-µm nominal pore-size black polycarbonate filters. After removing cellular material (0.2 µm filtration) and extracellular DNA with DNase I, we concentrated samples onto 0.02 µm Anodisc filters to estimate the abundance of free phage particles. Each phage-containing filter was then stained with SYBR Green I (Noble & Fuhrman 1998). We estimated population densities after capturing ten images from each filter with a CY3 filter set (ex: 550 nm, em: 570 nm) for *Synechococcus* or a FITC filter set (ex: 497 nm, em: 520 nm) for phage using a Zeiss microscope and image processing modules found in the base package of the Axiovision imaging software (Release 4.5 SP1).

We evaluated the influence of nutrient stoichiometry on community dynamics in three ways. First, we used repeated measures (RM) ANOVA to test for the effects of nutrient stoichiometry and time on *Synechococcus* and phage densities. Second, because cryptic dynamics can alter the temporal synchrony between hosts and parasites, we performed cross-correlation analyses for each chemostat using Auto Regressive Moving Average (ARMA) procedures (Lennon & Martiny 2008). In addition to determining whether populations in a chemostat positively or negatively covaried (i.e., in phase vs. out of phase, respectively) we used the cross-correlation analyses to test if the duration of time which viruses lagged behind host densities was altered by nutrient stoichiometry. Last, we estimated the effects of nutrient stoichiometry on the stability of microbial populations by calculating the inverse of the coefficient of variation (CV) over time.

### Coevolutionary dynamics

To test for the effects of nutrient stoichiometry on phenotypic coevolution, we tracked changes in infection patterns between *Synechococcus* and its phage over time. We isolated multiple (3-5) *Synechococcus* strains from each chemostat using serial dilution techniques six days prior to phage addition (day −6) and at days 9, 23, 72, 129, 148, and 166 after phage addition. We performed 10-fold dilutions of chemostat samples into 24-well plates containing sterile AN media that matched the nutrient supply ratio of the chemostat environment. When a dilute suspension of *Synechococcus* 7803 was incubated without shaking, single-colonies formed in the bottom of the wells (25 °C on a 14:10 light : dark cycle under 20 µE m^−2^ s^−1^). We picked these colonies with a Pasteur pipette and passaged them twice in 50 mL Erlenmeyer flasks using the same medium prior to harvesting. We then used epifluorescence microscopy to screen the cultures to make sure they did not contain free phage particles. Finally, these *Synechococcus* strains were concentrated *via* centrifugation, preserved in glycerol (10 % final concentration), and stored at −80 °C until reanimation for use in challenge assays, which are described below. We also isolated multiple (3-5) phage strains from the phage-amended chemostats on days 23, 72, 129, 148, and 166 through double plaque purification. This process involved adding 40 µL of 0.2 µm-filtered chemostat sample to a soft agar (0.8 % agar) overlay containing the ancestral *Synechococcus* WH7803 as the host. We then incubated the serially diluted plates (25 °C on a 14:10 light : dark cycle under 20 µE m^−2^ s^−1^), picked plaques that formed on the lawns of *Synechococcus*, and transferred to a log-phase batch culture of *Synechococcus* for propagation. The resulting phage lysates were syringe-filtered (1 µm) and preserved in glycerol (10 % final concentration) at −80 °C.

With the isolated chemostat strains, we quantified host resistance and phage infectivity using challenge assays. Each pair-wise challenge with strains isolated across chemostats within the same nutrient treatment was carried out in triplicate by adding 20 µL of a phage stock (∼10^7^ particles mL^−1^) to 200 µL of a dilute *Synechococcus* strain (∼10^6^ cells mL^−1^) in 96-well plates. Challenge assays were performed using the same medium from which the host strain was originally isolated (i.e., low vs. high N : P). Turbid cultures of *Synechococcus* WH7803 appear bright pink owing to the intracellular photosynthetic pigment phycoerythrin. When infected by SRIM8, however, cells are lysed and cultures become clear (Lennon & Martiny 2008). Based on this, we scored each *Synechococcus* strain as sensitive (and the phage strain as infective) if there was a lack of growth after a two-week incubation under continuous light (20 µE m^−2^ s^−1^) at 25 °C compared to control wells (n = 3) that contained heat-killed phage. To contend with the loss of a replicate chemostat in the N-limited treatment, we randomly selected *Synechococcus* strains across the two P-limited replicate chemostats to ensure that an identical number of strains were tested at each time point. In total, there were 18,050 pairwise challenges between chemostat-isolated strains of *Synechococcus* and phage that resulted in a bipartite infection matrix, which we used for all analyses related to phenotypic coevolution.

To quantify the effect of nutrient stoichiometry on coevolutionary dynamics, first, we used RM-ANOVA to test for trends in average resistance and average infectivity over the course of the chemostat experiment. Second, we used the infection matrix to perform time-shift analyses, which involved calculating the proportion of successful infections that occurred between hosts that were challenged against past, contemporary, and future phage strains (Gaba & Ebert 2009). We statistically analyzed the time-shift data using RM-ANOVA with nutrient treatment, phage treatment, and time as fixed effects, while the chemostat replicate identifier was treated as a random effect with a corARMA covariance matrix (Koskella 2014). To visualize the data, contemporary challenges were centered at a time-shift of zero, while interactions with past phage were represented in negative space and future interactions were represented in positive space. Last, to gain insight into potential mechanisms underlying stoichiometrically driven coevolution, we used community network analyses to calculate the connectance, nestedness, and modularity for each chemostat infection matrix. Connectance was calculated as the number of interactions divided by network size. We calculated nestedness using the NODF metric, which ranges from 0 (non-nested) to 1 (perfectly nested) and normalizes for matrix size. We used the LP-BRIM algorithm to find the partition that maximizes Barber’s modularity (Qb), which ranges from 0 (all interactions are between modules) to 1 (all interactions are within modules). The network statistics were calculated using 100,000 random Bernoulli simulations in the BiWeb package for MATLAB (Flores *et al*. 2011; Flores *et al*. 2016), http://github.com/tpoisot/BiWeb). We then tested for differences in connectance, nestedness, and modularity between the nutrient treatments using *t*-tests.

## RESULTS

### Nutrient limitation of *Synechococcus*

Prior to initiating the chemostat evolution trial, we conducted growth assays to test for N- and P-limitation of *Synechococcus*. After a period of acclimation in semi-continuous culture, the growth rates of N-limited *Synechococcus* increased by 87 % when additional N was supplied, but only by 35 % with the addition of P (*t*-test, *t*_8_ = 4.55, *P* < 0.001). In contrast, the growth rate of P-limited *Synechococcus* increased by 45 % with the addition of P, but only by 25 % with the addition of N (*t*-test, *t*_8_ = −3.36, *P* = 0.005). Results from this experiment suggest that while there may have been some degree of co-limitation, the two media sources used for subsequent evolution trials induced N- and P- limitation for *Synechococcus* (Fig. S1).

### Stoichiometry altered community dynamics

In the chemostat experiment, nutrient stoichiometry significantly affected the population dynamics of *Synechococcus* (RM-ANOVA; time × stoichiometry, *F*_62, 245_ = 2.43, *P* < 0.0001) and its phage (RM-ANOVA; time × stoichiometry, *F*_58, 230_ = 2.59, *P* < 0.0001). Under N-limitation, Synechococcus rapidly declined and reached its minimum mean density 35 days (range = 23 – 44) following phage addition (2 × 10^5^ ± 7 × 10^4^ cells mL^−1^, mean ± SEM, Fig. 1a). This decrease in host abundance corresponded with a peak in mean maximum density 25 days (range = 16 – 30) following phage addition (9 × 10^8^ ± 1 × 10^8^ particles mL^−1^, mean ± SEM). Over the next ∼50 days, the host densities slowly recovered, and entered a second phase of decline near day 100.

**Fig. 1.**
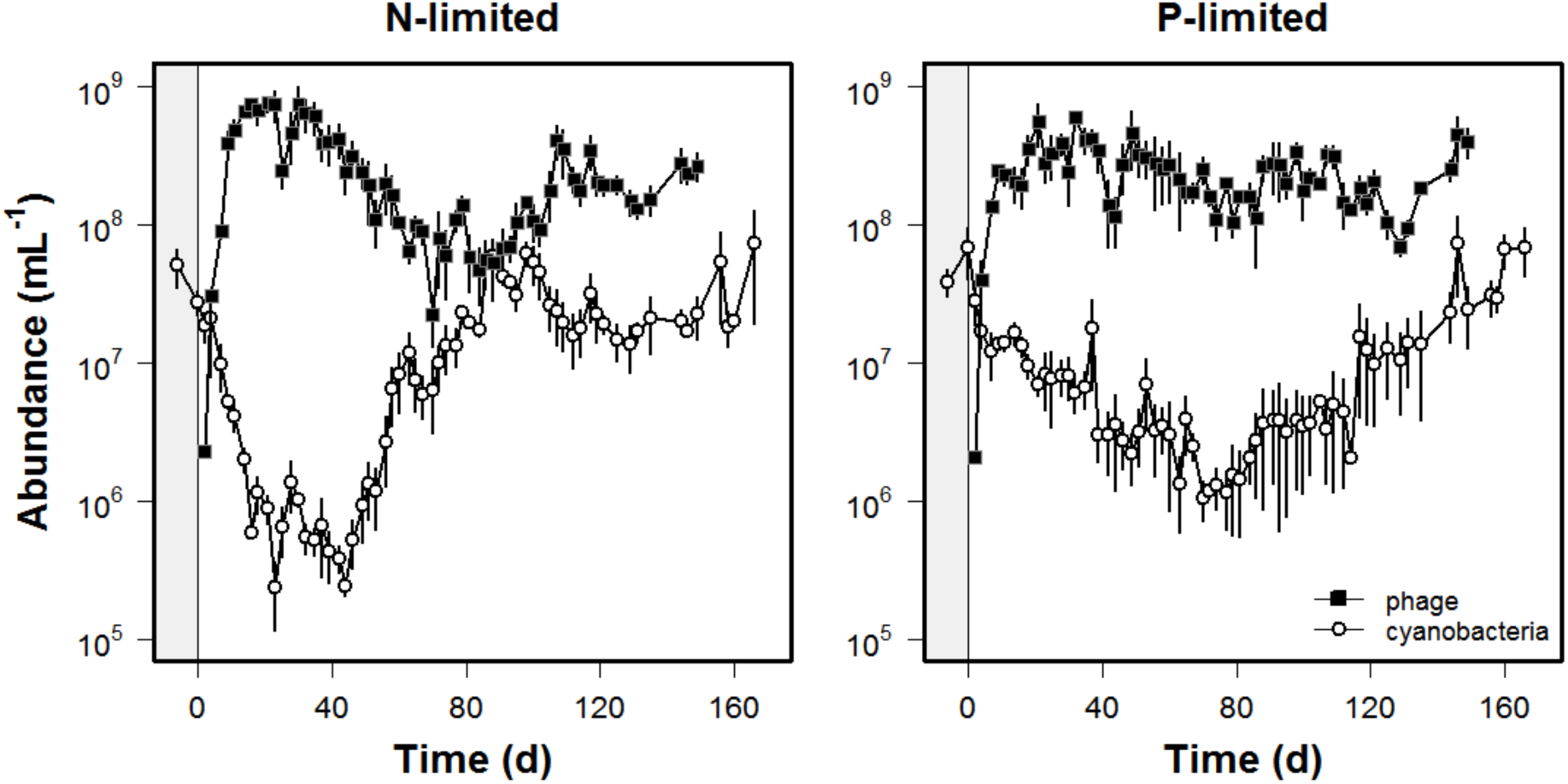
Microbial community dynamics were affected by nutrient stoichiometry. *Synechococcus* and phage densities were tracked in replicate (n = 3) chemostats receiving nitrogen (N)- or phosphorus (P)-limited media. Vertical lines at day 0 indicate time of phage amendment. See Fig. S2 for *Synechococcus* dynamics in the no-phage control chemostats. Data are represented as mean ± SEM.

*Synechococcus* and phage densities were significantly different and less dynamic under P-limitation (Fig. 1b). Hosts declined more slowly in P-limited chemostats following phage addition than in N-limited chemostats (*t*-test, *t*_4_ = −4.00, *P* = 0.02) and reached their mean minimum density 78 (range = 63 – 93) days following phage addition (4 × 10^5^ ± 6 × 10^4^ cells mL^−1^, mean ± SEM). Though hosts recovered, they did not reach pre-phage-addition abundances until the end of the experiment. Phage densities under P-limitation remained high and relatively constant (2 × 10^8^ ± 9 × 10^7^ particles mL^−1^, mean ± SEM) throughout the duration of the experiment. In the no-phage control chemostats, N- and P-limited *Synechococcus* densities remained constant over time (Fig. S2).

Nutrient stoichiometry also affected the temporal coherence and stability of *Synechococcus* and phage populations. After pre-whitening the time-series data using ARMA procedures, phage densities in N-limited chemostats were negatively correlated with host densities with lags ranging from zero to five days (*r* = −0.27 to −0.42, *P* = 0.009 – 0.063, Fig. S3). In contrast, under P-limitation, there was no correlation between phage and *Synechococcus* at any time lag (*r* = −0.20 to 0.22, *P* ≥ 0.23). Last, we found that cyanobacterial and phage densities were significantly more stable over time under P- than N-limited conditions (*Synechococcus*: *t*-test, *t*_4_ = 3.56, *P* = 0.024; *t*-test, phage: *t*_4_ = 3.83, *P* = 0.019). See Table S4 for summary statistics.

### Stoichiometry altered coevolution

Within nine days of phage addition, >50 % of the isolated *Synechococcus* strains were resistant to the ancestral phage in both the N- and P-limited chemostats (Fig. 2). Average resistance continued to increase over time (RM-ANOVA*, F*_6, 591_ = 14.2, *P* < 0.0001), but was not affected by stoichiometry (*F*_1, 4_ = 2.08, *P* = 0.22). The vast majority (97 %) of Synechococcus strains that were resistant to the ancestral phage were also resistant to all of the phage strains subsequently isolated from the chemostats. Phage-resistant *Synechococcus* did not revert to the sensitive phenotype when grown in the absence of phage consistent with the view that resistance is a heritable trait.

**Fig. 2.**
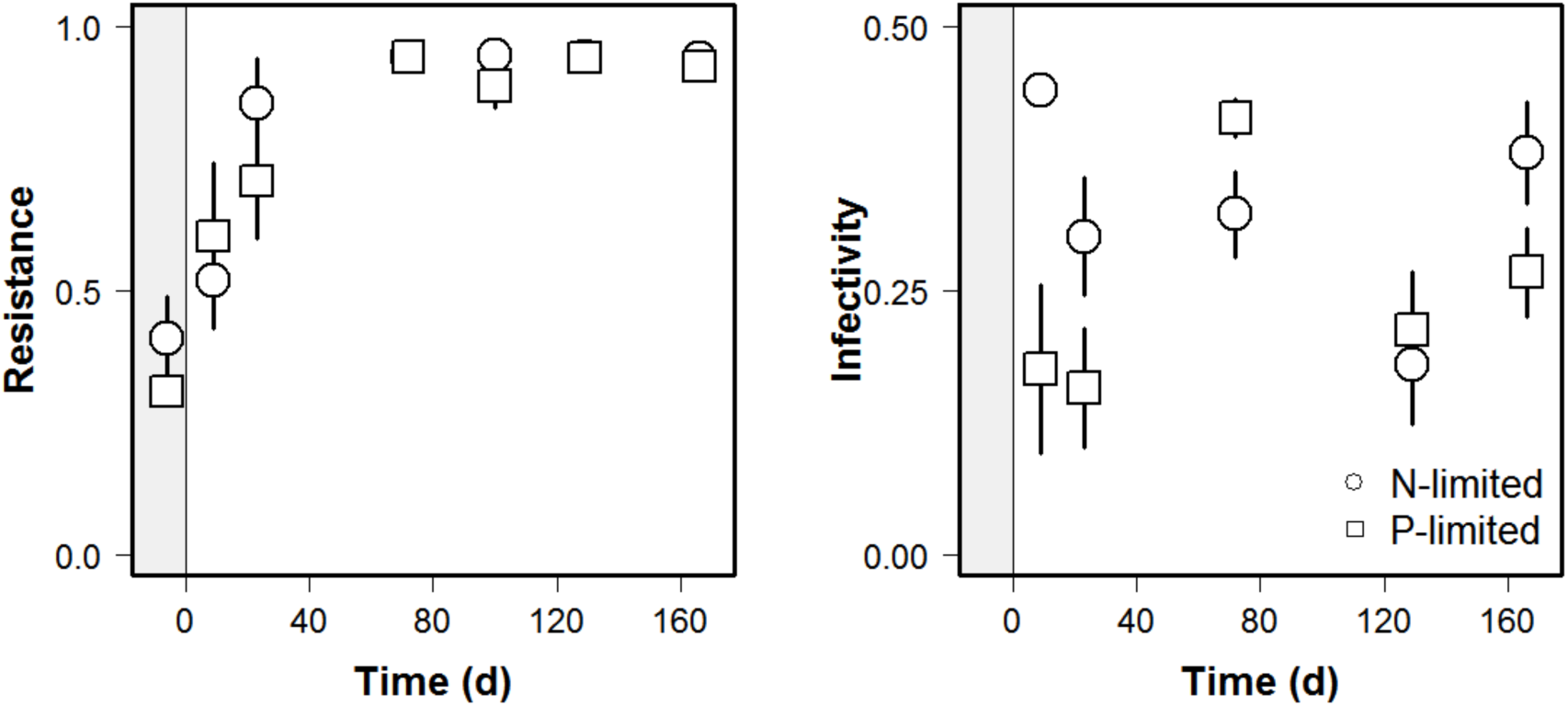
Average phage-resistance of *Synechococcus* strains increased monotonically over time under both nitrogen (N)- and phosphorus (P)-limitation (left). Resistance was calculated as the proportion of unsuccessful infections between all phage strains from the experiment when challenged against *Synechococcus* strains that had been isolated at a specific time point and is presented as the chemostat mean ± SEM. Average infectivity of phage over time (right) was calculated as the proportion of successful infections between all *Synechococcus* strains across the experiment when challenged against phage strains that had been isolated (on the ancestral host) at a specific time point and is presented as the chemostat mean ± SEM.

Phage infectivity also evolved, but in contrast to the host, was significantly affected by nutrient stoichiometry (Fig. 2; RM-ANOVA, stoichiometry × time: *F*_1, 601_ = 7.53, *P* = 0.006). In the no-phage control chemostats, *Synechococcus* strains remained susceptible to the ancestral phage. However, nearly all (∼ 96 %) of the derived phage from the phage-amended chemostats lost the ability to infect some of the *Synechococcus* hosts from the P-limited (n = 51) and N-limited (n = 22) no-phage chemostats. Despite this, we found evidence of host-range expansion. For example, a phage strain from a P-limited chemostat (day 129) was able to infect a *Synechococcus* strain (day 166) that was resistant to the ancestral phage. In addition, three phage strains from N-limited chemostats and two phage strains from P-limited chemostats (day 166) were able to infect phage-resistant *Synechococcus* isolated from other chemostats earlier in the study.

Time-shift experiments indicated that nutrient stoichiometry altered coevolutionary dynamics (Figs. 3 and 4). When hosts isolated from the phage-amended chemostats were challenged against current and past phage, interaction strengths were generally weak owing to the evolution of host resistance and phage infectivity (Fig. 3 and Fig 4 a, b; RM-ANOVA, stoichiometry × phage isolation time × bacterial isolation time, *F_1,597_* = 24.67, *P* < 0.0001). Hosts were more susceptible when challenged against future phage, but only for *Synechococcus* strains that were isolated earlier in the chemostat experiment (days −6, 9, and 23). The effect of nutrient stoichiometry was more evident in time-shift experiments where naïve hosts from the no-phage control chemostats were challenged against phage isolated from the phage-amended chemostats (stoichiometry × phage isolation time, F_1, 282_ = 7.25, *P* = 0.0075). Under these conditions, infectivity was not uniformly high across the time-shifts (Fig 3 and Fig. 4 c, d), as would be expected from purely arms-race dynamics. As a result, infectivity was 70 % greater on naïve hosts isolated from N-limited chemostats (0.61 ± 0.289) compared to naïve hosts isolated from P-limited chemostats (0.36 ± 0.332).

**Fig. 3.**
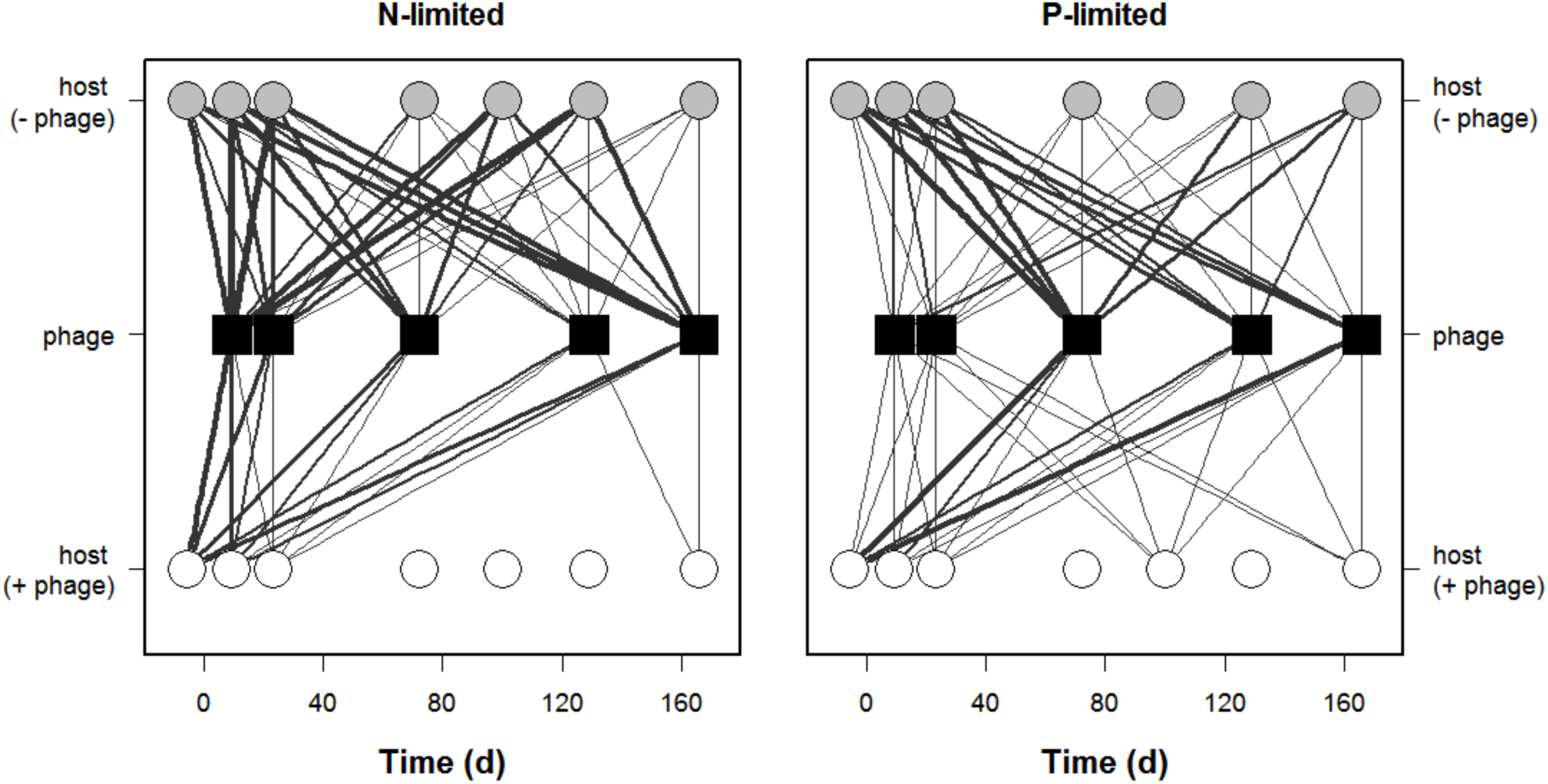
Phenotypic coevolution between hosts (*Synechococcus*) and phage was affected by nutrient stoichiometry. We calculated infectivity based on the proportion of successful infections between *Synechococcus* strains and phage strains that were isolated from chemostats at different time points. Infectivity is proportional to the width of the edges (lines) connecting nodes (symbols). Black squares correspond to phage isolated from the phage-amended chemostats, white circles correspond to *Synechococcu*s isolated from phage-amended chemostats, and grey circles correspond to naive *Synechococcus* isolated from no-phage control chemostats. The absence of a line indicates that *Synechococcus* isolates were resistant to a particular phage challenge.

**Fig. 4.**
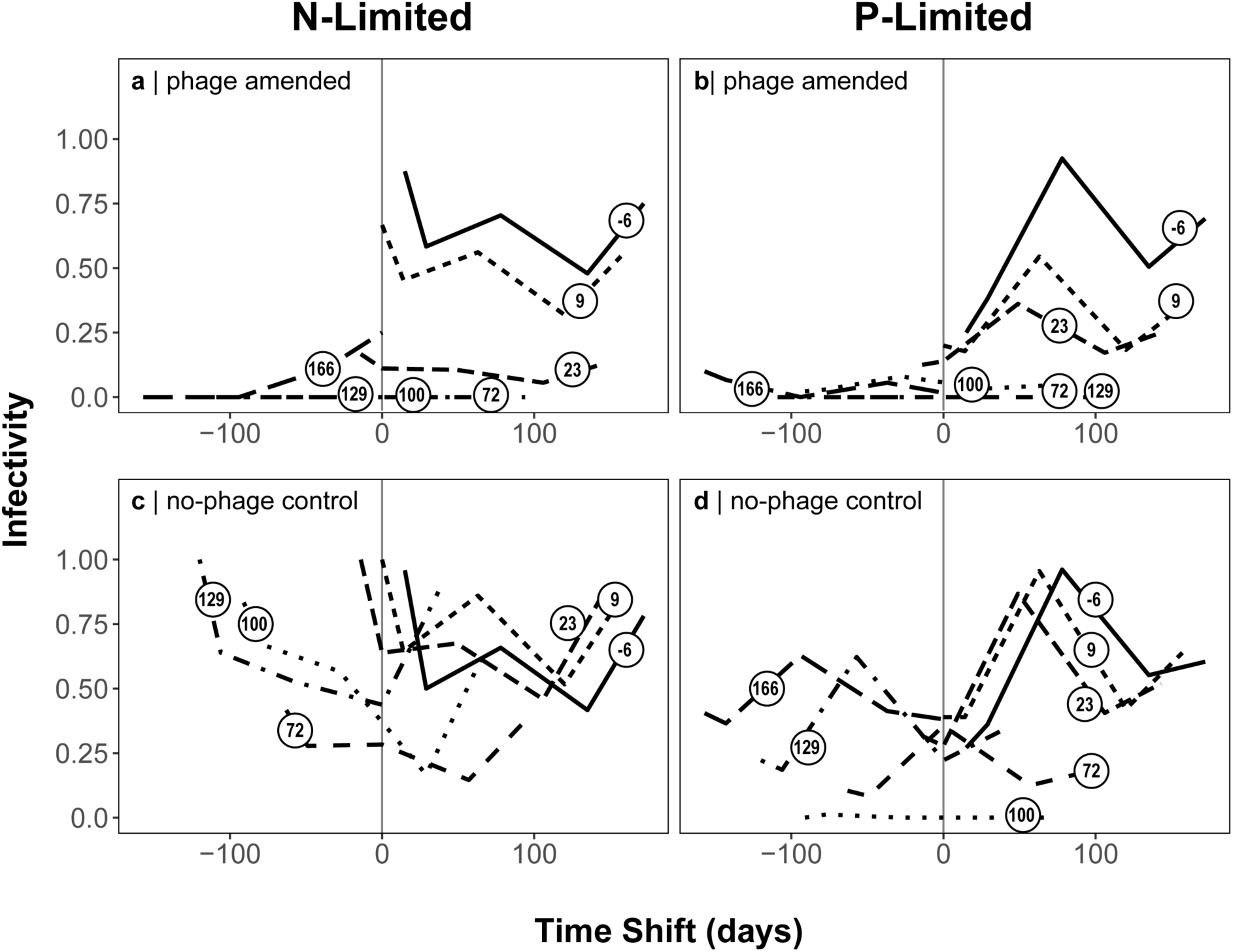
Time-shift analysis of host-phage infectivity reveals the effects of stoichiometry on coevolution. Contemporary interactions (i.e., those between a host and phage strain isolated at the same time point) are centered at time zero (grey vertical line) along the time-shift (horizontal) axis. Interactions with past phage are shifted to the left (negative values) and interactions with future phage are shifted to the right (positive values). Each black line corresponds with the mean infectivity for *Synechococcus* isolated from a specific time point as indicated by the open circle containing the isolation day (−6, 9, 23, 72, 100, 129, or 166). When comparing challenges between hosts and phage from phage-amended chemostats (a, b), infectivity was weak for hosts that were isolated after day 23 or when challenged against phage from the past owing to the evolution of resistance. Such findings are consistent with arms-race dynamics where directional selection gives rise to escalating host resistance. We also challenged phage against naive hosts from the no-phage control chemostats (c, d). From this, we found that infectivity was significantly higher under nitrogen (N)-limitation than phosphorus (P)-limitation, but overall, was lower than what would be expected under arms-race dynamics. Instead, the fluctuations in infectivity with respect to time-shift are consistent with negative frequency-dependent selection and reflect asymmetry in the coevolution between *Synechococcus* and phage.

Network analyses lent further support that nutrient stoichiometry affected host-phage coevolution. Patterns of infectivity were more nested and modular than expected by chance (*P <* 0.05), which is consistent with arms race dynamics and negative frequency-dependent selection, respectively. The infection networks from N- and P-limited environments had similar degrees of connectance (*t*_4_ = 1.47, *P* = 0.22) and nestedness (NODF, *t*_4_ = −0.37, *P* = 0.72). However, P-limited networks were 50 % more modular than N-limited networks (Fig. 5; *t*_4_ = −3.59, *P* = 0.02). See Table S6 for summary statistics.

**Fig. 5.**
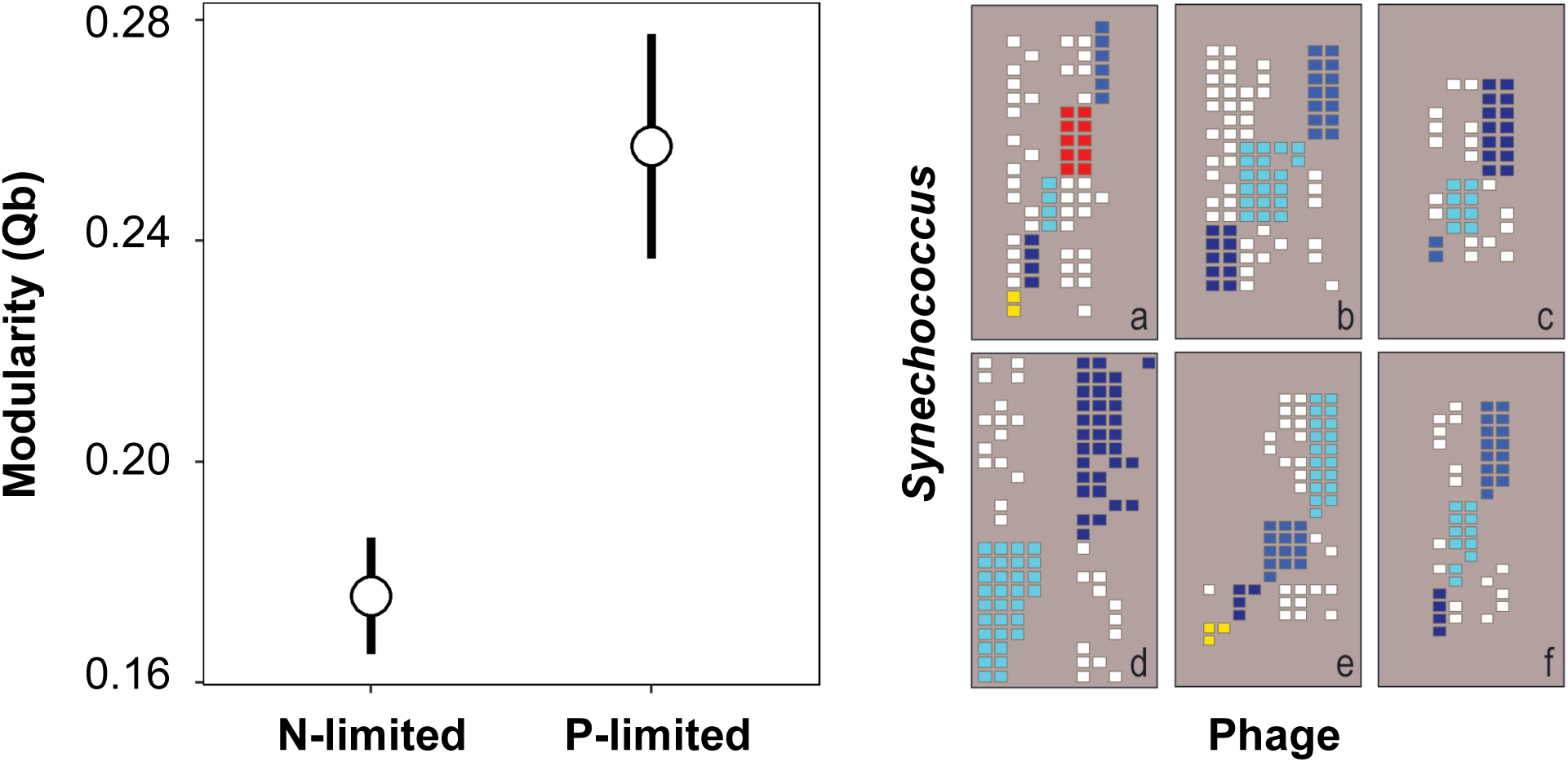
Host-phage infection networks based on interactions between *Synechococcus* and phage isolates that coevolved under nitrogen (N)- and phosphorus (P)-limitation. Networks were significantly nested, consistent with expectations of host-phage systems coevolving under arms race dynamics. However, the degree of nestedness was not affect by stoichiometry. Left panel: Networks were also significantly modular when compared to randomized networks, but the degree of modularity was significantly greater in P-limited networks. Right panel: Infection networks reflecting the modular structure of host-phage interactions within each chemostat. Each grouping of colored cells within a grey-colored background panel corresponds to a calculated module within the interaction network of N-limited (a – c) or P-limited (d – f) chemostats. That is, colored cells correspond to infectious interactions that could not be regrouped into a calculated module based on the network algorithm.

## DISCUSSION

The supply ratio of nitrogen (N) and phosphorus (P) had strong eco-evolutionary effects on marine *Synechococcus* and its phage. Communities were more stable under P-limitation than N-limitation, which likely was influenced by the nutritional flexibility and constraints of the host and phage populations, respectively. In addition, host-phage dynamics were influenced by the rapid invasion of resistant cyanobacteria which was followed by the appearance of evolved phage with altered host-ranges. Using time-shift assays and network analyses, we detected strong signatures of arms-race dynamics in both nutrient treatments. However, under P-limitation, we also observed patterns of increased network modularity consistent with negative frequency-dependent selection, which may reflect the underlying costs of maintaining defense and virulence traits in different nutrient environments. Our findings generate testable predictions regarding the mechanisms of local adaptation and coevolution for microorganisms among regions of the world ocean that are limited to varying degrees by N and P (Galbraith & Martiny 2015).

### Stoichiometry altered community dynamics

Consistent with predictions from the theory of ecological stoichiometry (Sterner & Elser 2002), we documented that *Synechococcus* and phage dynamics were highly sensitive to the N : P supply ratio. Under N-limitation, host-phage dynamics were less stable and out-of-phase, compared to P-limited chemostats (Fig. 1; Table S4). Historically, ecologists have attempted to explain such variation in populations and communities while assuming a single currency of resource. However, systems can exhibit a much wider range of behaviors when resource stoichiometry is explicitly considered owing to feedbacks that arise when interacting species vary in the degree to which their elemental composition is homeoestatically regulated (Andersen *et al*. 2004). For example, the molar N : P ratio of marine *Synechococcus* converges upon the canonical Redfield value of 16 : 1 in some environments (Garcia *et al*. 2016), but in other instances can exceed 100 : 1 (Bertilsson *et al*. 2003) reflecting the extreme plasticity of its biomass stoichiometry. In contrast, phage are thought to have a fixed elemental composition owing to their relatively simple structure, which minimally consists of genetic material (DNA or RNA) protected by a proteinaceous capsid (Jover *et al*. 2014). Biophysical models predict that the T4-like phage used in our study has a high P demand reflected by its low N : P ratio of approximately 7 : 1 (Jover *et al*. 2014). As a consequence, phage productivity is largely dependent on the P content of their hosts. For example, host lysis was delayed by 18 h leading to an 80 % reduction in phage burst size when P-limited *Synechococcus* WH7803 was infected with a myovirus (Wilson *et al*. 1996). Consistent with these reports, early in our experiment (day −6), phage infectivity was 50 % lower on P-limited vs. N-limited hosts (cf. Fig. 4A and 4B). It is possible, however, for phage to overcome the nutritional constraints of their microbial hosts. Many phage contain auxiliary metabolic genes that modify a host’s resource allocation (Monier *et al*. 2017). For example, phage-encoded genes for phosphorus acquisition (*pstS* and *phoA*) were upregulated when phosphorus-starved marine *Prochlorococcus* was infected by phage P-SSM2 (Zeng & Chisholm 2012). While the myovirus used in our study (S-RIM8) does not contain phosphate-binding protein or alkaline phosphatase genes, it does contain *phoH*. The exact function of *phoH* in marine microorganisms has not been demonstrated, but it is thought to encode a phosphate-starvation inducible protein that hydrolyzes ATP, thus liberating energy that can be used in P uptake (Goldsmith *et al*. 2011). Such findings suggest that nutrient stoichiometry can profoundly shape host-parasite dynamics (Aalto *et al*. 2015) and provide an ecological explanation for the patterns observed in our study (Fig. 1).

### Stoichiometry altered coevolution

Our results indicate that evolutionary processes were also important in creating feedbacks that influenced host-phage dynamics. Such feedbacks arise when rapid changes in host or parasite traits generate patterns that are not predicted from traditional ecological theory (Yoshida *et al*. 2007; Frickel *et al*. 2016). These so-called cryptic dynamics require the coexistence of multiple host genotypes, for example, through trade-offs in competitive ability and parasite defense (Yoshida *et al*. 2007). While there are many intracellular and extracellular mechanisms that bacteria can employ to resist phage infection, in *Synechococcus*, mutations in genes encoding for cell-surface receptors reduce or entirely eliminate attachment, thus precluding entry of the virus into the host (Stoddard *et al*. 2007; Marston *et al*. 2012). Selection for this phenotype could help explain the recovery of host densities in our chemostats (Fig. 1). However, resistance mutations in *Synechococcus* are often accompanied by a reduction in growth rate, the magnitude of which can vary depending on the identity of the host and virus (Lennon 2007). These fitness costs have important consequences for understanding community stability. Without a reduction in growth rate, resistant *Synechococcus* would outcompete the sensitive host and drive the phage population extinct. Moreover, fitness costs establish a trade-off that satisfies the requirement for cryptic dynamics to emerge (Yoshida *et al*. 2007). In sum, our data support the view that nutrient stoichiometry altered host-phage dynamics and stability via eco-evolutionary feedbacks (Lennon & Martiny 2008).

Nutrient stoichiometry had strong, yet importantly, asymmetric effects on microbial coevolution in our study. These effects were examined in light of the two primary modes by which antagonistic coevolution is thought to occur. The first is through arms-race dynamics involving gene-for-gene specificity, where directional selection leads to sweeps that are characterized by the escalation of resistance and infectivity. Arms-race dynamics were originally described for coevolving populations of plants and pathogens, but since then have been commonly reported in studies of bacteria and phage (Dennehy 2012; Koskella & Brockhurst 2014). The second mode of antagonistic coevolution involves negative frequency-dependent selection where parasites evolve to infect common hosts, which in turn favors rare host alleles. Negative frequency-dependent selection is often associated with infections that require matching alleles and is well documented in invertebrate systems (Decaestecker *et al*. 2007), but has also been described in some studies of bacteria and phage (Hall *et al*. 2011). While arms-race dynamics and negative frequency-dependent selection are often viewed as occupying different ends of the coevolutionary spectrum, they are not mutually exclusive (Agrawal & Lively 2002).

One powerful way to discern modes of coevolution is through the use of time-shift analyses. This approach involves determining the success of infections for combinations of hosts and parasites that are isolated from different time points in a controlled experiment or other longitudinal type of study (Gaba & Ebert 2009). When applied to challenge assays among host and phage isolates from our chemostats, time-shift analyses indicated that coevolutionary dynamics were significantly affected by nutrient stoichiometry (Fig. 3). The effect of nutrient stoichiometry was relatively weak when examining host strains isolated from phage-amended chemostats, which could reflect a mismatch between the time-shift interval and the rates of coevolution (Gandon *et al*. 2008). In contrast, the effect of nutrient stoichiometry was driven largely by interactions between evolved viruses that were challenged against *Synechococcus* isolated from no-phage control chemostats (Fig. 4 c, d). We expected that these naïve hosts would be uniformly and highly susceptible to all phage from the phage-amended chemostats. Instead, we found that infectivity fluctuated over time and was higher on N-limited vs. P-limited *Synechococcus*, reflecting expansion and contraction of the phage host-range (Fig. 4 c, d). Together, these findings are consistent with observations of asymmetric coevolution that have been attributed to the genetic constraints of viral populations (Lenski & Levin 1985). Asymmetry may also arise from the costs associated with being a generalist phage (Koskella & Brockhurst 2014), which may be dependent on the resource environment (e.g., nutrient stoichiometry). Such explanations have been used to explain host-phage coevolution where arms race dynamics give way to negative frequency dependent selection (Hall *et al*. 2011), in some cases depending on resource availability (Pascua *et al*. 2014).

The effects of nutrient stoichiometry on coevolution were further supported by results from our network analyses. With this approach, one can estimate the degree of nestedness that exists among pairs of hosts and phage relative to randomized data. Often, infection data from bacteria-phage systems are highly nested (Flores *et al*. 2011), which means they contain ordered subsets of phenotypes whereby hosts from later time points are resistant to earlier phages, and phage from later time points are able to infect earlier hosts (Weitz *et al*. 2013). In our study, infection networks were significantly nested, a pattern that arises from arms-race dynamics. However, the degree of nestedness was not affected by nutrient stoichiometry. Infection networks can also exhibit modularity, which is more consistent with negative frequency-dependent selection (Weitz *et al*. 2013). Modules reflect dense clusters of interacting host and phage compared to other strain combinations found in a bipartite infection matrix. Although less commonly documented, modularity has been observed in large-scale oceanic surveys of bacteria and phage, but has been attributed to local adaptation and the phylogenetic breadth of the hosts (Flores *et al*. 2013). However, theory suggests that modularity can emerge *via* coevolution between two populations at the local scale. Using a “relaxed lock and key” model, simultaneously nested and modular structures arose under simple chemostat conditions that assumed gene matching between phage tail-fibers and host receptors (Beckett & Williams 2013). Our results agree with these predictions, but suggest that network properties are affected by host nutrition. Specifically, we found that host-phage interactions were 50 % more modular under P-limitation than N-limitation, suggesting that nutrient stoichiometry constrains host-phage interactions leading to increased specialization. Such findings are consistent with the view that resources can influence the modes of coevolution based in part on the fitness costs associated with host defense and infection strategies (Pascua *et al*. 2014)

Mechanistically, there are many ways that nutrient stoichiometry could shape bacteria-phage coevolution. For example, the “dangerous nutrients” hypothesis predicts that trajectories of coevolution can be influenced when receptors used by nutrient-limited hosts also serve as the targets of phage adsorption (Menge & Weitz 2009). However, it does not appear that myoviruses, including the strain used in this study, attach to the protein receptors of cyanobacteria that are used for nutrient transport. Instead, evidence from whole-genome sequencing of the host used in our study suggests that phage-resistant *Synechococcus* accumulate mutations in hypervariable genomic islands that encode for lipopolysaccharide (LPS) (Marston *et al*. 2012), a major component of the outer membrane in Gram-negative bacteria. The structural complexity of LPS is affected by nutrient limitation (Brelles-Marino & Boiardi 1997) and such changes in molecular structure of the cell membrane can interfere with phage adsorption leading to resistance (Leon & Bastias 2015). Nevertheless, modification of tail-fibers allow phage to overcome resistance of marine cyanobacteria in some instances (Schwartz & Lindell 2017) while the acquisition of host-like P-assimilation genes may aid in the successful infection of nutrient-limited hosts (Kelly *et al*. 2013).

## Conclusions

Most organisms live in environments where they are limited by the relative or absolute amount of one or more essential resources. It is well established that such variation in nutrient stoichiometry can regulate ecological phenomena ranging from species interactions to ecosystem-level processes. We demonstrated that nutrient stoichiometry also affects the evolutionary dynamics of microbial communities likely through differences in the expansions and contraction of virus host-ranges under N- vs. P-limited conditions. Identifying the targets of selection in contrasting nutrient environments will help elucidate the genetic mechanisms of coevolution and the trade-offs associated with defense and virulence traits. While our study offers promising avenues to better understand bacteria-phage interactions in the aquatic environments, nutrient stoichiometry is also important in determining the nutrition, health, and disease susceptibility for non-microbial hosts and their microbiomes(Smith & Holt 1996), which has implications for understanding the persistence, emergence, and evolution of infectious diseases. In the oceans, phage represent a significant source of mortality for marine cyanobacteria like *Synechococcus*, which play a central role in the regulation of biogeochemical processes, including energy flow, carbon sequestration, and the cycling of nitrogen (N) and phosphorus (P). Our findings suggest that the coevolutionary process was differentially affected by the availability of N and P in ways that could influence the ecology and evolution of one of the most abundant and functionally important groups of microorganisms on Earth.

## Acknowledgments

We acknowledge technical support from R Morrison, M Carroll, and BK Lehmkuhl; ancestral strains from MF Marston; discussion with CM Lively; feedback from BK Whitaker, KJ Locey, WR Shoemaker, NI Wisnoski, V Kuo, ME Muscarella, DA Schwartz, RZ Moger-Reischer, JM Palange on earlier versions of the manuscript. Financial support was provided by the National Science Foundation (0851143, 0851113) and the US Army Research Office Grant W911NF-14-1-0411. We dedicate this work to the memory of VH Smith.

**Author contributions**
MLL, SWW, and JTL designed study; MLL performed research; MLL and JTL analyzed data; MLL, SWW, and JTL wrote paper.

