## Supplementary Materials for "Nutrient stoichiometry shapes microbial coevolution"

**Table S1** Chemostats were supplied with a modified version of artificial seawater medium (AN) (Waterbury, J.B. & Willey 1988) that induced nitrogen (N) or phosphorus (P) limitation (see Fig. S3) while maintaining similar equilibrium densities of *Synechococcus* (see Fig. S4). We autoclaved a solution containing dissolved Base Salts in 500 mL of Nanopure water. Once cool, we added 0.1-μm sterile-filtered Nutrient Solutions, cycloheximide (a eukaryotic inhibitor), the Trace Metals Solution (Table S2), and Vitamin Solution for enrichment and purifications (Table S3). Finally, we brought the mixture to 1 L final volume with autoclaved Nanopure water.

| **Base Salts** | **Mass** (g $L^{-1}$) |
| --- | --- |
| NaCl | 21.61 g |
| $MgSO_4 \cdot 7H_2O$ | 7.39 g |
| $MgCl_2 \cdot 6H_2O$ | 4.07 g |
| $CaCl_2 \cdot 2H_2O$ | 1.47 g |
| KCl | 0.75 g |

| **Nutrient Solutions** | **Stock Concentration** (g $L^{-1}$) | **Volume** (mL $L^{-1}$) | |
| --- | --- | --- | --- |
| | | N-limited | P-limited |
| EDTA (disodium salt) | 1.00 | 5.00 | |
| $Na_2CO_3 \cdot H_2O$ | 3.00 | 5.00 | |
| $NaHCO_3$ | 6.10 | 5.00 | |
| $NaNO_3$ | 4.00 | 6.26 | 12.52 |
| $K_2HPO_4$ | 8.40 | 0.63 | 0.32 |
| **Trace Metals Solution** | | 1.00 | |
| **Vitamin Solution** | | 1.00 | |
| **Cycloheximide** | 50 mg $L^{-1}$ | 100 µL | |

**Table S2** Recipe for Trace Element Solution used in the modified AN media. All trace metals were dissolved into 1 L of Nanopure water then filter (0.1-μm) sterilized.

| **Compound** | **g L$^{-1}$** |
|---|---|
| $ZnSO_4 \cdot 7H_2O$ | 0.222 |
| $MnCl_2 \cdot 4H_2O$ | 1.4 |
| $Co(NO_3)_2 \cdot 6H_2O$ | 0.025 |
| $Na_2MoO_4 \cdot 2H_2O$ | 0.39 |
| $C_6H_8O_7 \cdot H_2O$ | 6.25 |
| $C_6H_8FeNO_7$ | 6.0 |

**Table S3** Recipe for Vitamin Solution used in the modified AN medium, which was used in the medium for enrichment and purification of *Synechococcus* strains in this study. Each compound was prepared as a neat stock (unless otherwise noted) then combined in the volumes listed below to prepare a 100 mL solution. The final solution was filter sterilized (0.1-µm) and stored at -20° C in 50 mL aliquots.

| **Compound** | **Stock Concentration** ($g\ L^{-1}$) | **Amount per 100 mL** |
|---|---|---|
| Inositol | | 100 mg |
| Thiamine · HCl | | 20 mg |
| Vitamin $B_{12}$ | 1.0 | 0.1 mL |
| Biotin | 0.1 | 1 mL |
| Folic Acid | 2.0 | 0.1 mL |
| p-aminobenzoic acid | 2.0 | 0.5 mL |
| Niacin (Nicotinic acid) | 1.0 | 10 mL |
| Ca d-pantothenate | 2.0 | 10 mL |
| Pyridoxine | 1.0 | 10 mL |
| Nanopure water | | To 100 mL |

**Table S4** Summary statistics for *Synechococcus* (Syn) and phage in experimental chemostats. Nutrient limitation refers to the nitrogen (N) : phosphorus (P) supply ratio where N-limited chemostats received medium with a 10 : 1 N : P ratio and P-limited chemostats received medium with a 40 : 1 N : P ratio. Each experimental unit was assigned a unique Chemostat ID. Half of the chemostats phage amended with S-RIM8 (Phage Treatment = +) while the remaining chemostats served as no-phage controls (Phage Treatments = -). Equilibrium abundances of *Synechococcus* and phage were estimated as the mean ± SEM of densities over the time series based on methods described in the main text. Mean stability was measured for *Synechococcus* and phage within a chemostat as the inverse of the coefficient of variation of population densities over time.

| Type | Nutrient Limitation | Phage Treatment | Mean Density ± SEM | | | | Stability | |
|---|---|---|---|---|---|---|---|---|
| | | | Syn | | phage | | Syn | phage |
| Treatment averages | N | + | 1.7E+07 | ± 1.31E+07 | 2.5E+08 | ± 1.39E+08 | 0.75 | 1.05 |
| | P | + | 1.1E+07 | ± 1.09E+07 | 2.4E+08 | ± 9.49E+07 | 0.59 | 1.45 |
| | N | - | 1.3E+07 | ± 3.69E+06 | · | · | 2.05 | · |
| | P | - | 1.8E+07 | ± 9.02E+06 | · | · | 1.14 | · |

| Type | Nutrient Treatment | Chemostat ID | Phage Treatment | Mean Density ± SEM | | | | Stability | |
|---|---|---|---|---|---|---|---|---|---|
| | | | | Syn | | phage | | Syn | phage |
| Phage-amended | N | N2 | + | 1.8E+07 | ± 1.21E+07 | 1.5E+08 | ± 7.92E+07 | 0.84 | 1.11 |
| | N | N3 | + | 1.6E+07 | ± 1.00E+07 | 2.9E+08 | ± 1.46E+08 | 0.93 | 1.16 |
| | N | N5 | + | 1.7E+07 | ± 1.65E+07 | 3.1E+08 | ± 1.61E+08 | 0.59 | 1.12 |
| | P | P2 | + | 1.3E+07 | ± 1.43E+07 | 1.6E+08 | ± 5.91E+07 | 0.52 | 1.58 |
| | P | P4 | + | 1.1E+07 | ± 6.66E+06 | 2.4E+08 | ± 8.84E+07 | 0.94 | 1.59 |
| | P | P5 | + | 9.4E+06 | ± 1.04E+07 | 3.1E+08 | ± 1.11E+08 | 0.52 | 1.63 |
| No-phage Control | N | N1 | - | 1.3E+07 | ± 3.69E+06 | · | · | 2.05 | · |
| | P | P1 | - | 2.5E+07 | ± 1.03E+07 | · | · | 1.43 | · |
| | P | P3 | - | 9.9E+06 | ±4.02E+06) | · | · | 1.42 | · |

**Table S5** Coevolutionary dynamics in phage-bacteria infection matrices have been inferred from network metrics including modularity and nestedness. We calculated network statistics based on the matrix of infection data resulting from pairwise challenges of phage and *Synechococcus* that were isolated from N- and P-limited chemostats using the BiWeb program in Matlab (Flores *et al.* 2011), available at https://github.com/tpoisot/BiWeb. Network size is calculated from the number of interactions and reflected the number of strains that were isolated from a chemostat. Connectance, the proportion of possible links between strains, was calculated as the number of interactions divided by network size. Barber's Modularity (Qb) was calculated for each chemostat system using the LP-BRIM algorithm to find the partition that best maximized modularity within a matrix. Nestedness (NODF) is based on overlap and decreasing fill, which returns a value ranging between 0 and 1 (where 1 indicates a perfectly nested structure) and normalizes for matrix size, allowing for differing sizes of matrices to be compared. All calculations were based on 100,000 random Bernoulli simulations as described in the main text.

| **Treatment** | **Chemostat ID** | **Size** | **Connectance** | **Modularity** ($Q_b$) | **Nestedness** (NODF) |
|---|---|---|---|---|---|
| NL | N2 | 144 | 0.257 | 0.162 | 0.331 |
| | N3 | 126 | 0.194 | 0.197 | 0.374 |
| | N5 | 65 | 0.210 | 0.168 | 0.242 |
| PL | P2 | 253 | 0.220 | 0.216 | 0.387 |
| | P4 | 180 | 0.168 | 0.276 | 0.266 |
| | P5 | 95 | 0.144 | 0.279 | 0.224 |

**Figure S1** Results from nutrient limitation assay. After acclimating *Synechococcus* in, we found that the change in *Synechococcus* growth rate in batch culture significantly increased in response to the addition of limiting resource (N or P) following acclimation to N-limited (N : P = 10 : 1) or P-limited (N : P = 40 : 1) conditions. Data are represented as mean ± SEM (n = 5).

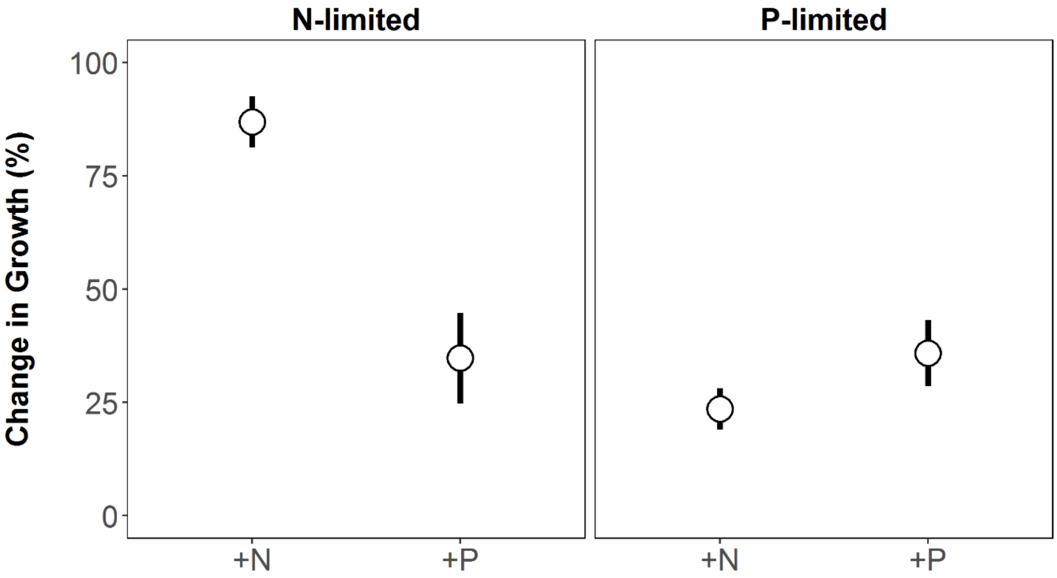

**Figure S2** Population dynamics of *Synechococcus* WH7803 in chemostats were relatively stable over time in the absence of phage (*i.e.*, no-phage controls). The N-limited treatment (left) only had one chemostat following the loss of the second control chemostat due to contamination. There were two replicates in the no-phage control P-limited chemostats (data represent mean ± range). Vertical line corresponds to *Synechococcus* densities prior to phage introduction in the other phage-amended chemostats.

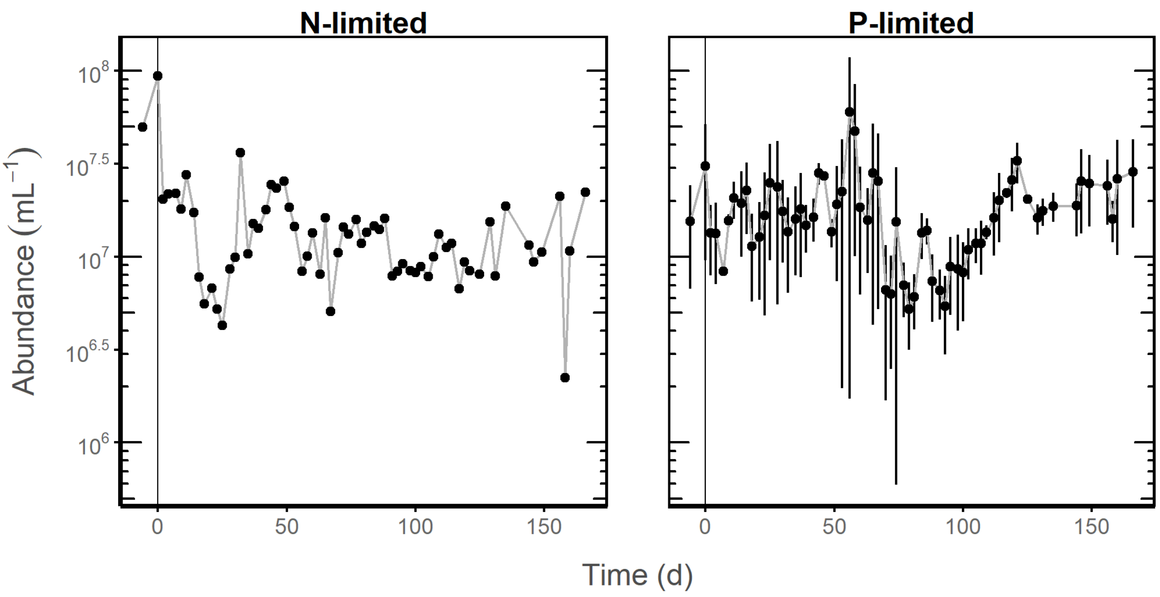

**Figure S3** Temporal coherence of *Synechococcus* and phage population densities under nitrogen-limited (left) or phosphorus-limited conditions (n = 3). We pre-whitened the log-$_{10}$ transformed time series using Autoregressive Moving Average (ARMA) before calculating cross-correlation coefficients (CCF). Results from CCF can range from -1 where host-phage dynamics are out-of-phase to +1 where host-phage dynamics are in-phase. The lag corresponds to the amount of time (in units of sampling dates) that the *Synechococcus* and phage densities were shifted when calculating the CCF. Negative time lags suggest that *Synechococcus* densities were correlated with phage densities from the past, while positive lags suggest that phage densities track *Synechococcus* densities from the past. The blue shaded region in each plot represent two standard-error limit after white noise correction.

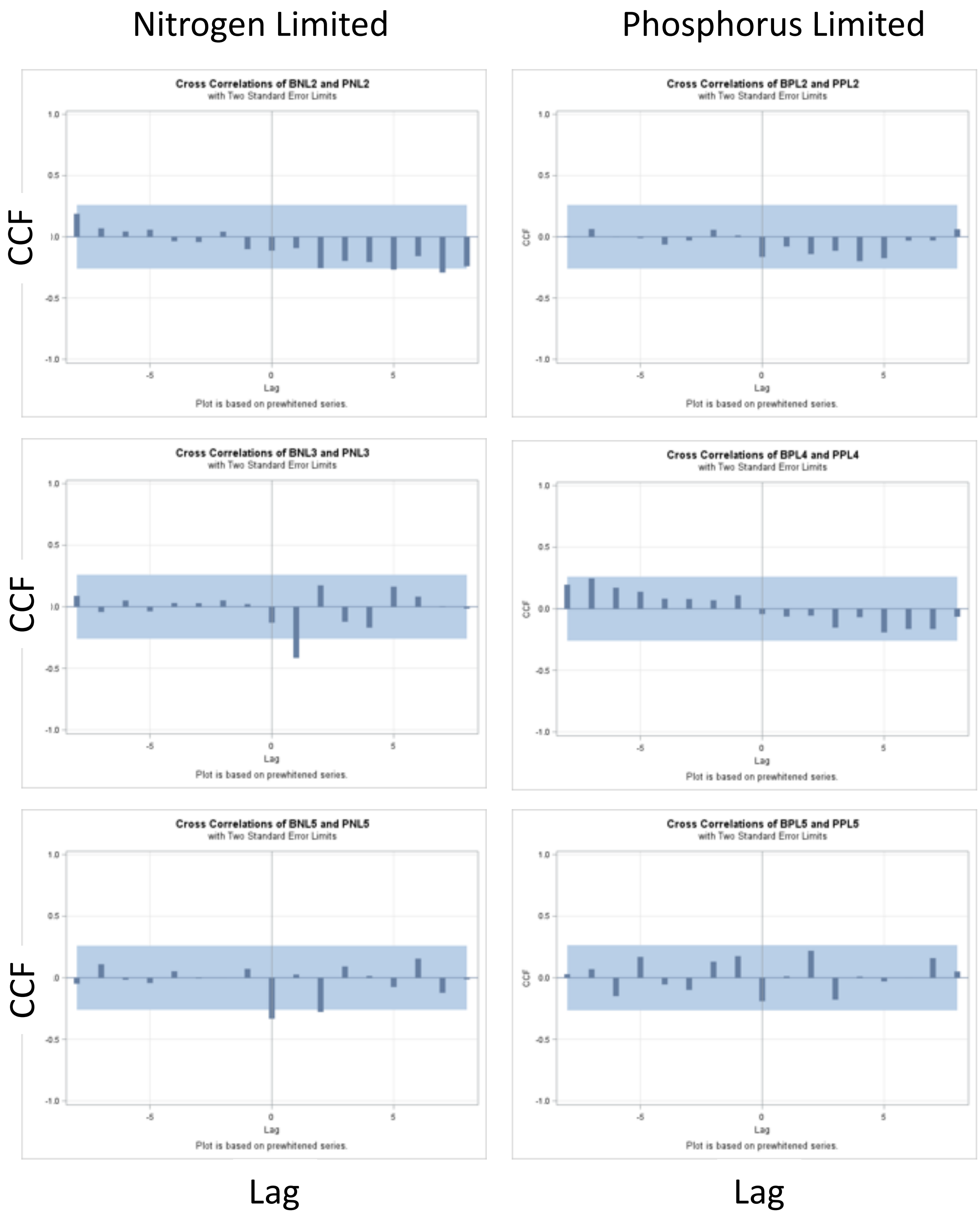
Nitrogen Limited
Phosphorus Limited
Cross Correlations of BNL2 and PNL2
with Two Standard Error Limits
Cross Correlations of BPL2 and PPL2
with Two Standard Error Limits
Cross Correlations of BNL3 and PNL3
with Two Standard Error Limits
Cross Correlations of BPL4 and PPL4
with Two Standard Error Limits
Cross Correlations of BNL5 and PNL5
with Two Standard Error Limits
Cross Correlations of BPL5 and PPL5
with Two Standard Error Limits
CCF
Lag
Plot is based on prewhitened series.

**Figure S4** Infection matrices for N-limited (a, b, c) and P-limited (d, e, f) replicate chemostats. Cyanobacteria and phage interactions within a chemostat were quantified using triplicate challenge assays with 3-5 *Synechococcus* and 3-5 phage strains isolated per tested time-point (n = 7). Most of the successful lytic infections between a *Synechococcus* (column) and phage (row) strain were constrained between strains isolated early in the experiment (prior to day 23, white cells). Phage-resistant *Synechococcus* phenotypes appeared quickly within the chemostats and persisted throughout the duration of the experiment (blue cells). Axes correspond to the number of strains within the rows or columns.

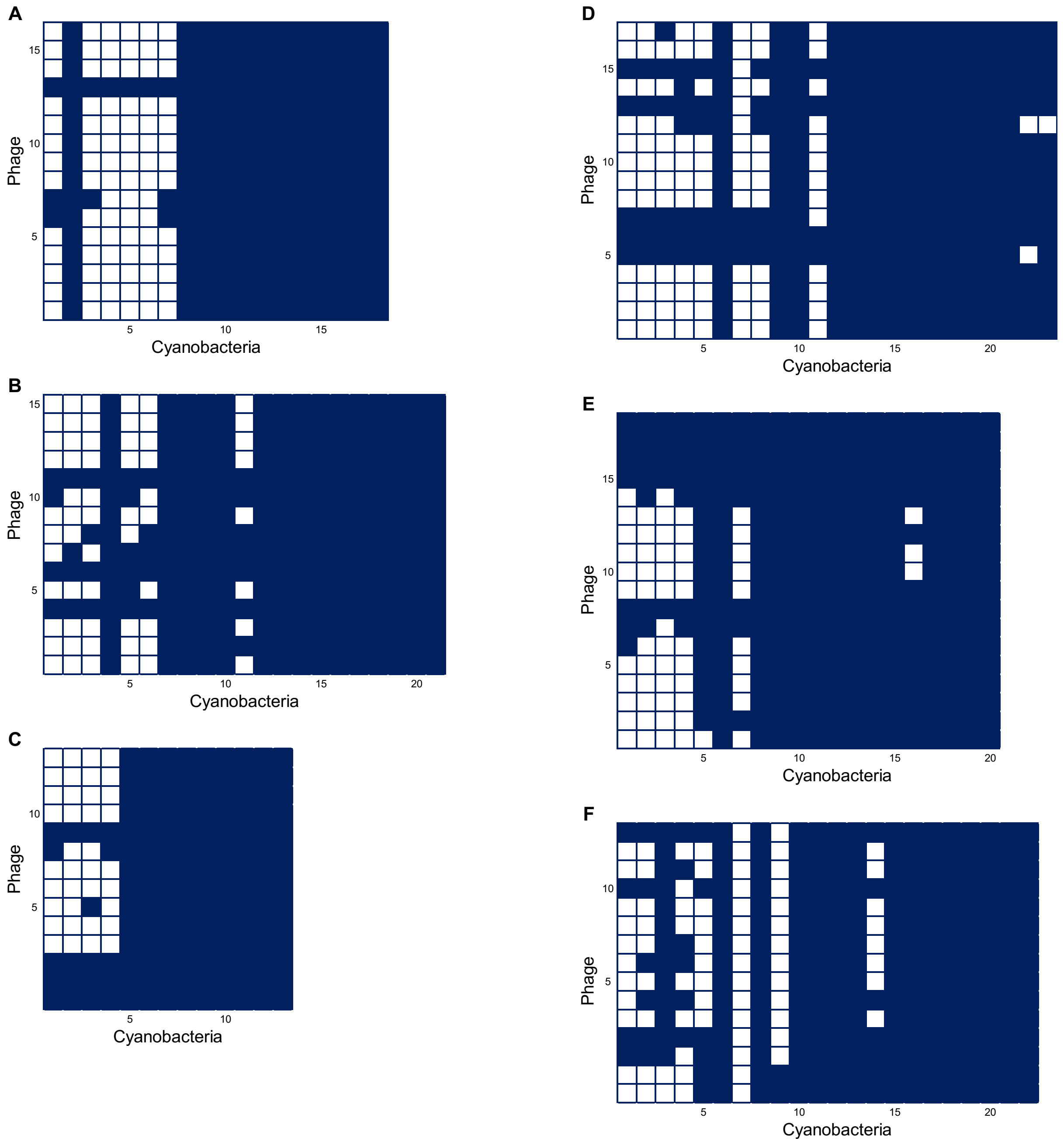
A
B
C
D
E
F
Phage
Cyanobacteria
5
10
15
20
